# Two interdigitated fine-scale channels for encoding motion and stereopsis within the human magnocellular stream

**DOI:** 10.1101/2022.04.20.488911

**Authors:** B. Kennedy, P. Bex, D.G. Hunter, S. Nasr

## Abstract

In humans and non-human primates (NHPs), motion and stereopsis are processed within fine-scale cortical sites, including V2 thick stripes and their extensions into areas V3 and V3A that are believed to be under the influence of magnocellular stream. However, in both species, the functional organization (overlapping vs. interdigitated) of these sites remains unclear. Using high-resolution functional MRI (fMRI), we found evidence for two interdigitated channels within human extrastriate areas that contribute to processing motion and stereopsis. Across multiple experiments that included different stimuli (random dots, gratings, and natural scenes), the functional selectivity of these channels for motion vs. stereopsis remained consistent. Furthermore, an analysis of resting state functional connectivity revealed stronger functional connectivity within the two channels rather than between them. This finding provides a new perspective toward the mesoscale organization of the magnocellular stream within the human extrastriate visual cortex, beyond our previous understanding based on animal models.

---

In the visual cortex of humans and NHPs, the magnocellular processing stream consists of multiple fine-scale cortical sites that are distributed across extrastriate visual cortex ^1, 2^. In both species, these cortical sites are involved in processing motion and stereopsis (depth), two important but fundamentally different aspects of any visual stimuli. To this day, in either species, it remains unknown whether motion and stereopsis are encoded within two separate (interdigitated) sites within the magnocellular stream or if the same (overlapping) sites contribute to encoding both features.

In NHPs, stimulus motion and stereopsis are processed selectively within V2 thick stripes ^3-10^. The information encoded within these stripes is further processed within areas V3, V3A and MT ^1, 11^. Stereo-selective neuronal columns (and clusters) have been reported within all these areas ^12-18^. Motion-selective neuronal columns are also found frequently within areas V3 ^12, 19^, V3A ^20-22^ (but see also ^23, 24^) and MT ^25-27^. However, the relative organization of motion-and stereo-selective sites within these areas remains largely unknown.

With recent advances in high-resolution neuroimaging techniques, homologous columnar organizations have been shown in human visual cortex. Specifically, multiple studies have shown evidence for cortical columns across areas V2, V3 and V3A that contribute to encoding stereopsis ^28-30^ and motion ^31, 32^. Similar to NHPs, and based on their stimulus selectivity and functional connections, these cortical columns appear to be under the influence of magnocellular stream ^2, 31^. Here again, the relative organization of motion-and stereo-selective sites within the early visual areas remains largely unknown due to technical challenges in functional imaging of such small brain sites.

Thus, despite the abundance of evidence for existence of motion-and stereo-selective neuron columns and clusters across areas V2, V3 and V3A in humans and NHPs, the detailed nature of their organization (overlapping vs. interdigitated) remains unknown. Consequently, it is not clear whether motion processing is conducted in parallel with that of stereopsis or if the processing of motion is intermixed with that for stereopsis. This lack of knowledge has limited our ability to infer the underlying circuitry at mesoscale levels.

Here, we tested the hypothesis that motion and stereopsis are processed through interdigitated channels within the human extrastriate visual areas. Using high-resolution fMRI, we localized the fine-scale motion-and stereo-selective sites across areas V2, V3 and V3A in the same individuals. Using this technique, we also tested the functional properties of these sites, including their contribution in orientation, spatial frequency (SF), motion direction and depth encoding plus their functional connections during resting state. The result of these tests revealed that motion-and stereo-selective sites are interdigitated within several cortical areas, including V2, V3 and V3A, forming two channels within the magnocellularly influenced regions.

## Results

Fifteen human subjects (6 females; aged 23–44 years old) with normal vision participated in this study (see Online Methods and Table S1). Each subject participated in multiple experiments, all using high-resolution fMRI (voxel size = 1 mm isotropic). Experiment 1 localized motion-and stereo-selective sites in different individuals. Experiment 2 independently confirmed that these sites are mainly under the influence of the magnocellular stream based on their interdigitated organization relative to color-selective sites (Experiment 2a), their orientation-sensitivity (Experiment 2b), and their low-SF preference (Experiment 2c). Experiments 3 and 4 confirmed that the organization of these sites is largely preserved irrespective of the stimulus shape. Experiment 5 tested the selective contribution of these sites to motion-direction and depth encoding. Finally, Experiment 6 tested whether sites with similar stimulus selectivity are functionally interconnected bases on an analysis of resting-state functional connectivity.

### Experiment 1. Motion and stereopsis are encoded selectively within interdigitated sites

In all participants (*n=*15), we first localized the cortical sites in areas V2, V3 and V3A that responded selectively to moving (compared to stationary) stimuli (Figure 1A) and to depth-varying (compared to depth-constant) random dot stereograms (RDS; Figure 1B). For each individual, the border of retinotopic visual areas was defined during a separate scan session, based on independent stimuli ^33^.

**Figure 1.**
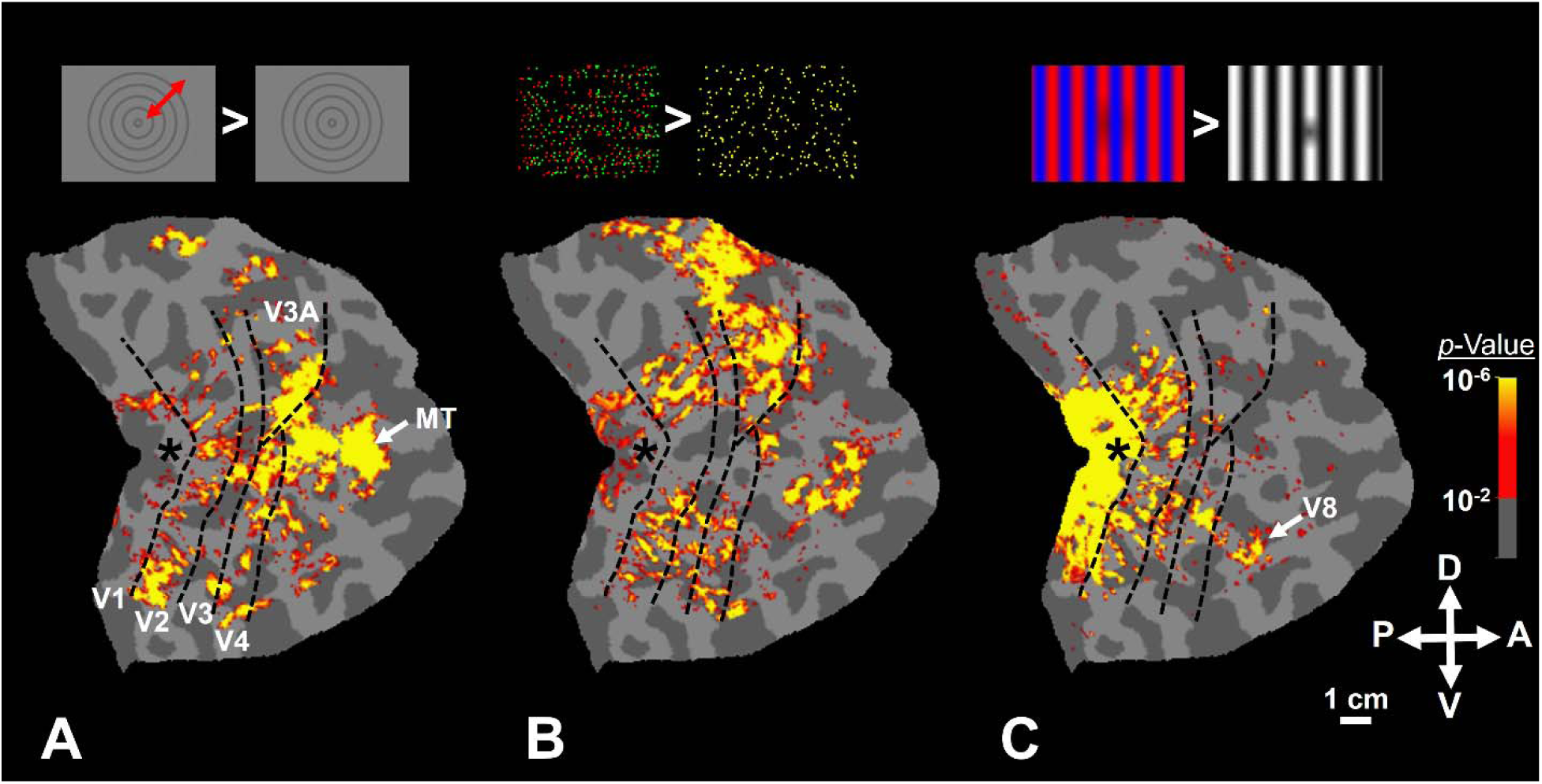
Motion, stereopsis and color evoke selective activity within the extrastriate cortex of one individual subject, overlaid on the subject’s flattened cortex (Experiments 1 and 2a). Panels show the selective activity evoked by contrasting the response to moving vs. stationary (**Panel A**, 3D vs. 2D RDS (**Panel B**) and color-vs. luminance-varying stimuli (**Panel C**; Experiment 2a). A schematic representation of the stimuli is also illustrated on top of each panel (see Online Methods). In all experiments, we detected stripy activity sites, starting at V1-V2 border, and extending into areas V3 and V4. The stripy/patchy pattern of activity in areas V2 and V3 is consistent with the expected shape of stripes reported in histological studies (Tootell and Taylor, 1995; Adams et al., 2007). As reported previously (Tootell and Nasr, 2017), color-selective sites were rarely detected within area V3A. In each panel, the border of retinotopic visual areas is shown with black dashed lines. The asterisks show the foveal representation in area V1. The location area MT (**Panel A**) and V8/VO (**Panel C**) are indicated by white arrows.

As demonstrated in Figure 1 for one individual subject, motion- (Figure 1a) and stereo-selective (Figure 1b) activity was often topographically elongated in a direction perpendicular to the V1-V2 border, extending through areas V3 and V3A, consistent with the known topography of stripes in humans ^34, 35^. In a few subjects in which the field-of-view extended far enough (e.g. Figure 1a), we were also able to identify the motion-selective area MT, near the posterior end of the medial temporal sulcus ^36^.

When co-registered, motion-and stereo-selective sites formed an interdigitated pattern (Figures 2 and 3 and Table S2) with only minor spatial overlap (≤5.89 ± 4.46%) (See Online Methods). In areas V2, V3 and V3A, this level of overlap was significantly smaller than chance level (t(14)>2.15, *p*<0.05), defined as when the selectivity maps were spatially shuffled. This result provides the first direct evidence for interdigitation of mesoscale motion-and stereo-selective sites across human visual areas V2, V3 and V3A.

**Figure 2.**
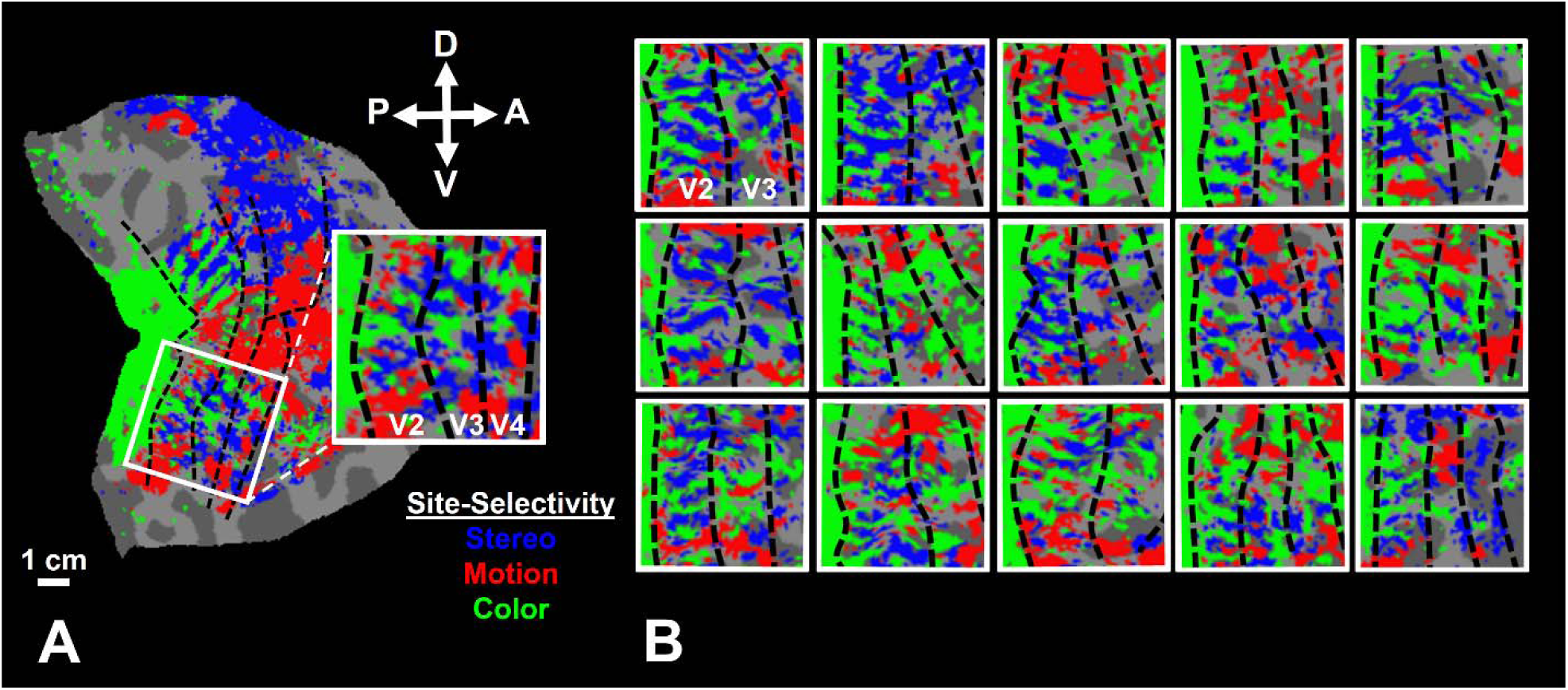
The co-localization of motion-, stereo-and color-selective sites (Experiments 1 and 2a). **Panel A** shows the data from the same individual as in Figure 1. **Panel B** shows the interdigitation of motion-, stereo-, and color-selective sites in 15 hemispheres other than the one showed in **Panel A**. In all hemispheres, the selective sites are distributed in areas V2 and V3 and formed an interdigitated pattern. The between-subjects-variability in the spatial organization of these sites is apparent. In all panels, the overlapping regions are excluded from the maps.

**Figure 3.**
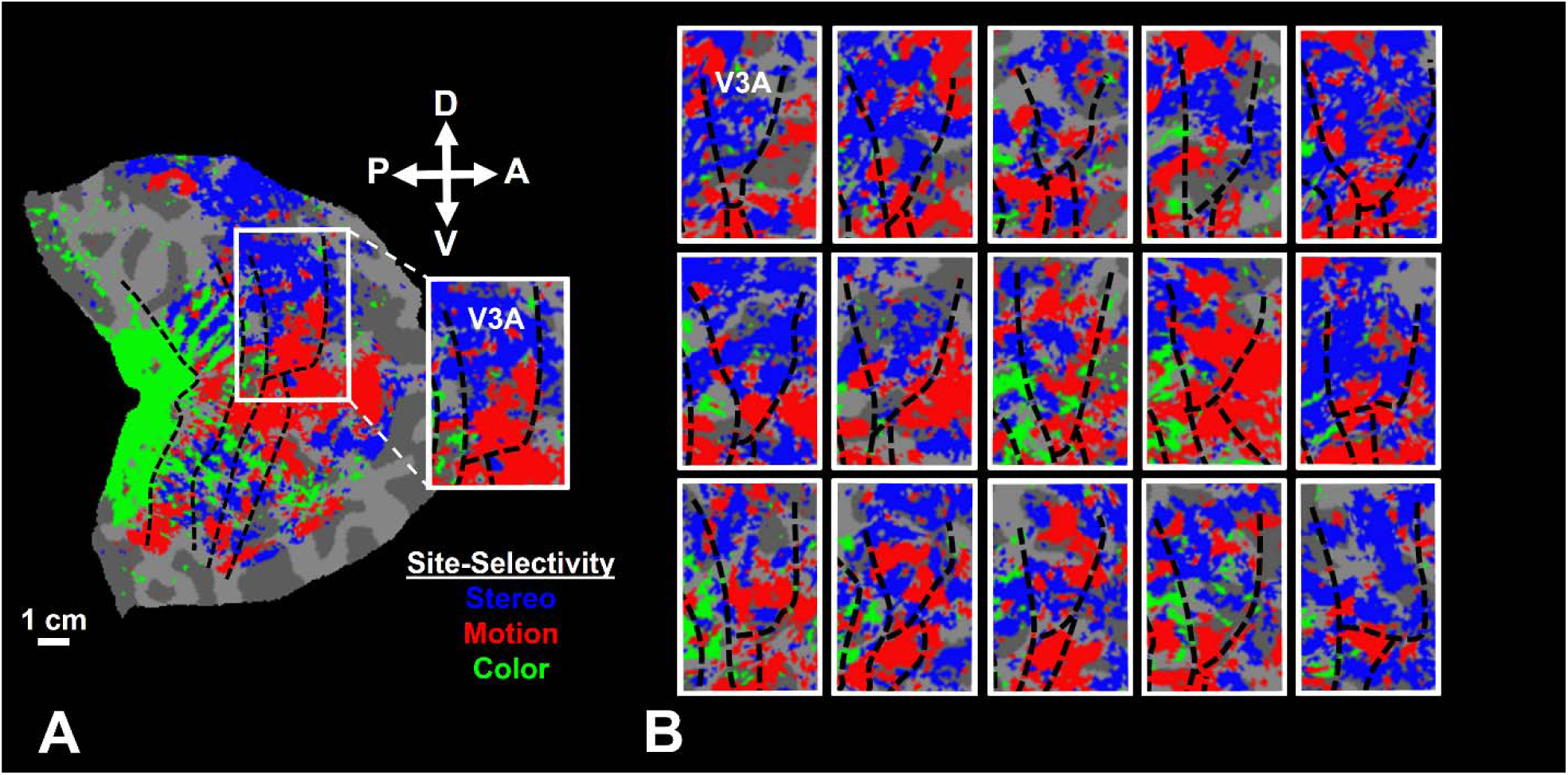
The co-localization of motion-, stereo-and color-selective sites in area V3A (Experiments 1 and 2a). **Panel A** inset shows area V3A from the same individual as in Figures 1. **Panel B** shows the same data from 15 other hemispheres. In all hemispheres, interdigitated motion and stereo-selective sites are apparent in V3A, while color-selective sites are rarely detected in this area. Here again, the between-subjects-variability in in the exact location of these sites is apparent. Other details are the same as **Figure 2**.

### Experiment 2. Motion and stereopsis are encoded within the magnocellularly-influenced sites

We tested the hypothesis that motion-and stereo-selective sites are localized within those fine-scale cortical regions that are under the influence of the magnocellular stream. If true, we expected these sites: *(i)* to interdigitate with color-selective sites that comprise the parvocellular stream (Experiment 2a), *(ii)* to show stronger orientation selectivity compared to color-selective sites (Experiment 2b) and *(iii)* to show a relatively higher preference for lower SF compared to higher SF stimuli (Experiment 2c).

### Experiment 2a. Motion-and stereo-selective sites were localized outside color-selective sites

Based on studies in NHP, we expected magnocellularly influenced sites to show no overlap with color-selective sites that comprise the parvocellular stream ^10, 37, 38^. To test this expectation, Experiment 2a localized color-selective sites based on their stronger response to isoluminant color-varying vs. luminance-varying stimuli in our participants (*n=*15; Table S1), as in prior tests of color selectivity ^28, 39^. Then the location of those color-selective stripes was compared to the location of motion-and stereo-selective sites.

Results of this test showed that, beyond area V1, color-selective sites formed a striped topography beginning at the V1-V2 border, extending through area V3 ^28^ (Figure 1C). In a few subjects for whom the field-of-view included the fusiform area (e.g. Figure 1C), presumptive area V8/VO could also be identified in the posterior portion of middle fusiform sulcus ^39-41^. However, in contrast to motion-and stereo-selective sites, color-selective sites were rarely detected within area V3A (Figures 1C and 3 and Table S2).

Moreover, as expected from the suggested segregation of magnocellular and parvocellular streams ^10, 37, 38^, when co-registered (Figure 2), we found minimal overlap between motion-and stereo-selective activity maps relative to color-selective maps (≤4.71 ± 1.85%) (Table S2). Across areas V2 and V3, the level of this overlap was significantly below chance level (t(14)>4.94, *p*<10^−3^), defined as when the selectivity maps were spatially shuffled. These results indicated that motion-and stereo-selective sites are preferentially located outside color-selective sites that comprise the parvocellular stream.

### Experiment 2b. Motion-and stereo-selective sites show stronger orientation selectivity compared to color-selective sites

In NHPs, magnocellularly influenced thick stripes in V2 and their extension in other areas could be detected reliably based on their stronger orientation selectivity relative to parvocellularly influenced thin stripes ^3, 4, 42-45^. Accordingly, we next measured the level of orientation selectivity of motion-, stereo-and color-selective sites across areas V2 and V3. Area V3A was excluded from this test because color-selective sites were rarely detected in this area (see above).

When subjects (*n=*15) were presented with orientation-varying stimuli (see Online Methods), motion-and stereo-selective sites showed a significantly stronger orientation-selective response compared to color-selective sites across areas V2 (t(14)=3.56; *p<*0.01) and V3 (t(14)=2.63; *p*=0.02) (Figure S1). The same analysis did not yield any significant difference between the level of orientation-selective activity measured within motion-vs. stereo-selective sites (t(14)<1.81; *p*>0.10). These results support the hypothesis that motion-and stereo-selective sites overlap the magnocellularly influenced thick stripes in V2 and their likely extension in V3.

### Experiment 2c. Motion-and stereo-selective sites show stronger preference for lower SF compared to color-selective sites

Magnocellularly influenced regions are expected to show a stronger preference for lower rather than higher SF stimulus components ^46, 47^. Experiment 2c compared the SF preference of motion-, stereo-and color-selective sites. When subjects (*n=*11; Table S1) were presented with gratings with different SFs (0.1 to 5.79 c/deg; see Online Methods), we found a significant SF × site-type interaction on activity evoked within areas V2 (*p*<10^−6^) and V3 (*p*<10^−4^) (see also Table S3). In both areas, motion-and stereo-selective sites showed a stronger response to gratings of lower SF (peak ≤0.27 c/deg) compared to higher SF (Figure S2). In contrast, color-selective sites showed a significantly stronger response to higher SF (peak ∼0.73 c/deg) gratings. This result is consistent with the hypothesis that the magnocellular stream has a stronger influence on motion-and stereo-selective compared to color-selective sites. Here again, area V3A was excluded from this test since it shows little or no color selectivity (see above).

### Experiment 3. Variations in stimulus structure (shape) did not affect the interdigitated organization of motion-and stereo-selective sites

In Experiment 1 we localized motion-and stereo-selective sites using gratings and RDS, respectively. The SF and orientation content of gratings (barrow band) and RDS (broad band) significantly differ. To test whether the interdigitated organization observed in Experiment 1 were related to differences in motion and stereo, rather than differences in stimulus bandwidth, in Experiment 3 we tested whether (or not) the selectivity of activity maps changed in response to different “carrier” stimuli.

### Experiments 3a and 3b. Use of random dots rather than gratings to activate motion-selective sites

Two independent experiments were conducted to test the effect of stimulus shape on the level of motion selectivity. In Experiments 3a, one group of subjects (*n=*11; Table S1) was presented with random dots that, across different blocks, were either stationary or moving <u>radially</u> (see Online Methods). In Experiment 3b, another group of subjects (*n=*11; Table S1) was presented with dots that were either stationary or moving <u>translationally</u>. As demonstrated in Figures 4A and S3, the overall pattern of motion-selective activity remained similar between Experiments 1 and 3. As demonstrated in Figures 4B, in both experiments, moving dots evoked a significantly stronger selectivity within the motion-selective sites (defined based on moving vs. stationary gratings) compared to the stereo-selective sites (*p*<10^−3^; see also Tables S4 and S5). These results indicate that motion-selective sites across V2, V3 and V3A areas respond selectively to motion irrespective of the carrier stimuli.

**Figure 4.**
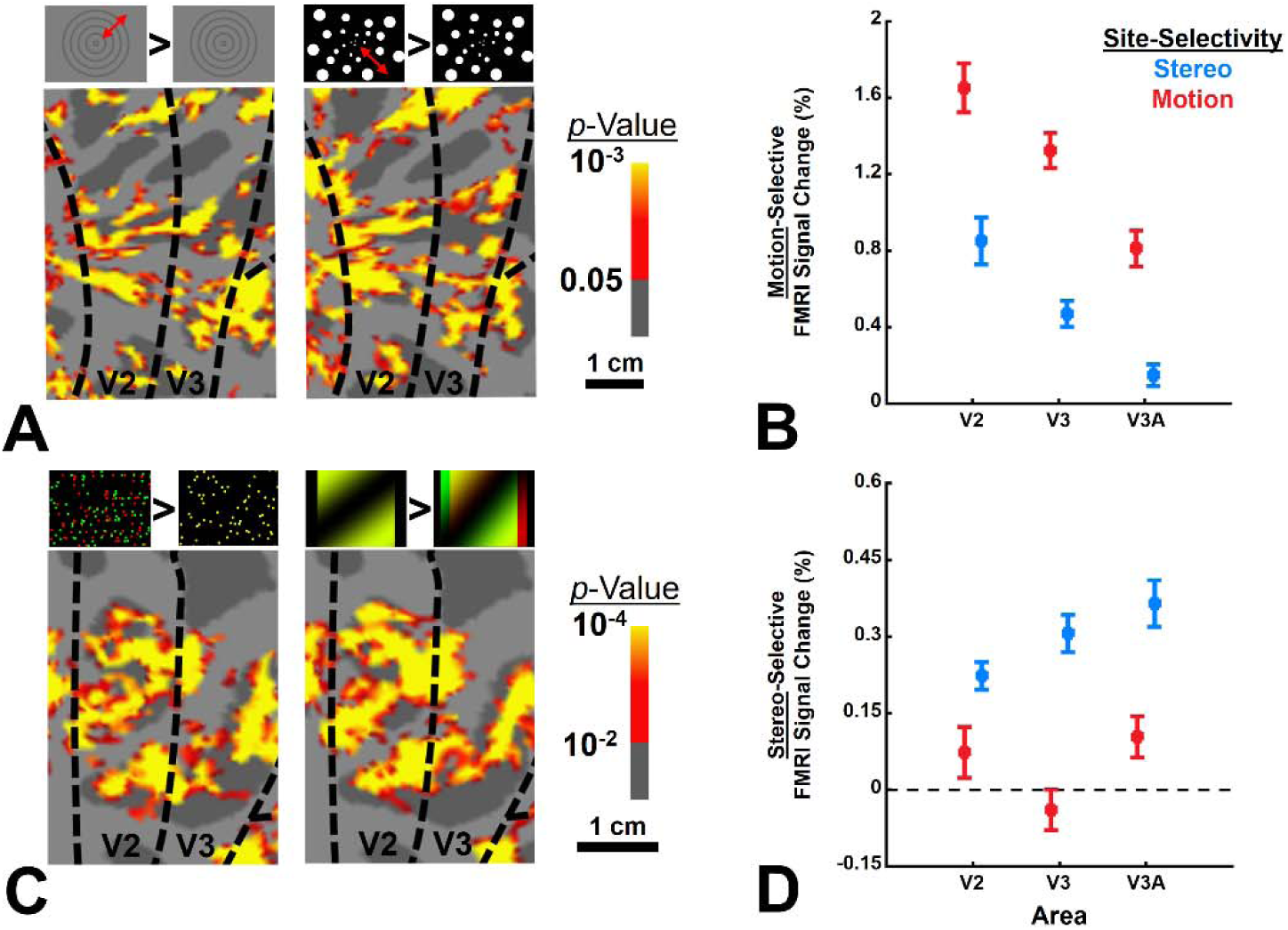
The effect of changing the shape of stimuli (from gratings to random-dots and vice versa) on the pattern of activity evoked within motion-and stereo-selective sites (Experiment 3). **Panels A and C** show the motion and stereo-selective activity, respectively. For each map, the type of stimuli used in the experiment is demonstrated on top of the panel. **Panels B and D** show the level of motion-and stereo-selective activity across areas V2, V3 and V3A, evoked by random-dots and gratings, respectively. The regions of interest (motion-vs. stereo-selective) in each area were localized independently based on gratings and random-dots. Consistent with activity maps, the level of evoked activity by ‘moving vs. stationary’ and ‘3D vs. 2D’ stimuli was higher in motion-and stereo-selective sites, respectively. It suggests that changing the shape of stimuli did not alter the overall pattern of activity evoked within motion-and stereo-selective sites. Error bars show one standard error of mean.

### Experiment 3c. Use of gratings rather than RDS to activate stereo-selective sites

To test whether depth-varying stimuli other than RDS could also evoke a selective response within the stereo-selective sites, a group of subjects (*n=*11; Table S1) were presented with depth-varying (3D) vs. depth-invariant (2D) gratings (SF=0.25 c/deg) (see Online Methods). Here again, the pattern of stereo-selective response remained similar to that observed in Experiment 1 (Figure 4C). As demonstrated in Figure 4D, depth-varying stimuli evoked a significantly stronger response within the stereo-selective sites (defined based on RDS) compared to the motion-selective sites (*p*<10^−3^; see also Tables S6). These results indicate that stereo-selective sites across V2, V3 and V3A areas respond selectively to the stimulus binocular disparity irrespective of its spectral content. Together, the results of Experiment 3 suggest that the interdigitated organization of motion-and stereo-selective sites remains consistent, irrespective of the carrier stimuli (see also Experiment 4).

### Experiment 4. Stereo-selective sites could be localized without disparity “oscillation”

The RDS stimuli used in Experiments 1 and 3 oscillated in depth (see Online Methods) to enhance 3D perception ^48^. This approach raised the possibility that the current interdigitated organization of motion-vs. stereo-selective sites reflects separated representation of two axes of motion (i.e. motion-in-depth vs. motion-in-fronto-parallel-planes). Previous studies in NHPs ^49^ and humans ^50^ showed that MT activity evoked by motion-in-depth can be differentiated from activity evoked by motion-in-fronto-parallel plane. However, the contribution of earlier visual areas to this process remains unclear. Accordingly, Experiment 4 tests whether the interdigitated organization of motion-vs. stereo-selective sites across area V2, V3 and V3A (as shown in Experiments 1-3), represents two separate axis of motion (i.e. motion-in-depth vs. motion-in-fronto-parallel-planes).

A group of subjects (*n=*6; Table S1) were scanned as they were presented with non-oscillating 3D vs. 2D natural scenes (see Online Methods). If the stereo-selective sites were localized based on their selectivity for motion-in-depth (rather than stereopsis per se), we expected them to lose their selectivity in this test. Alternatively, if those sites were selectively responding to the stimulus stereopsis irrespective of its oscillation, we expected the overall activity maps would remain similar to those generated based on RDS. The results of this test showed strong stereo-selective activity evoked by non-oscillating natural scenes (Figure S4). Importantly, the spatial distribution of stereo-selective sites across areas V2, V3 and V3A remained similar between Experiments 1 and 4 (Figure S4A) and the level of stereo-selective activity (Figure S4B) was significantly stronger within stereo-selective sites (as localized independently based on RDS) rather than motion-selective sites (*p*<0.01; see also Tables S7). These results rule out the possibility that the interdigitated stereo-and motion-selective sites represent two axes of motion. Moreover, they further support the hypothesis that the interdigitated organization of motion-and stereo-selective sites does not depend on the stimulus spatial structure (see also Experiment 3).

### Experiment 5. Selective contribution to direction and depth discrimination

Motion-selective sites are expected to be selectively involved in discriminating stimuli based on their motion direction ^4, 32, 51^. In Experiment 3 (see above), subjects were presented with radial (i.e. centrifugal vs. centripetal) and translational (i.e. leftward vs. rightward vs. upward vs. downward) motion (Online Methods). Here, we compared the motion direction sensitivity of evoked response within motion-and stereo-selective sites during those tests. For radial motion, direction sensitivity was defined as the absolute difference between the responses evoked by centripetal vs. centrifugal motion. For translational motion, direction sensitivity was averaged over all opponent pairs of motion directions.

In both tests (Figure 5A and S4), we found a significantly stronger direction sensitivity in the motion-compared to the stereo-selective sites (*p*<0.01; see also Tables S8 and S9). The overall level of direction sensitivity weakened significantly from areas V2 to V3A (*p*<10^−3^), perhaps due to the simplicity of the motion stimulus used in these tests. In any event, these results support the hypothesis that, compared to stereo-selective sites, motion-selective sites contributed more strongly in signaling variation in motion direction.

**Figure 5.**
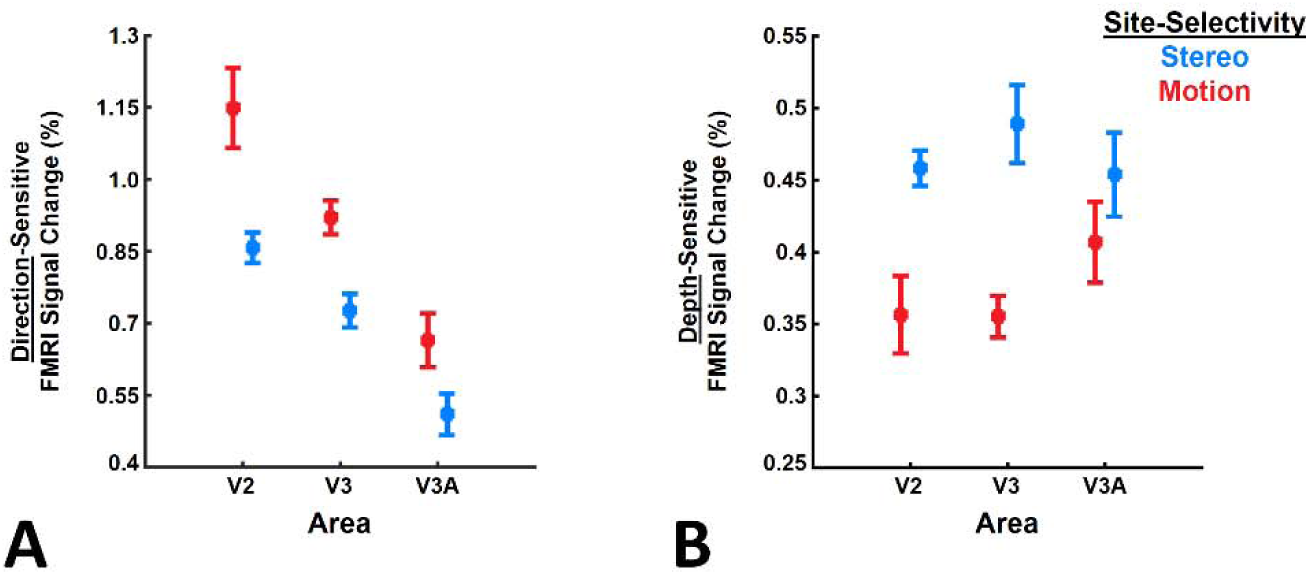
Sensitivity to the stimulus motion direction and depth measured across motion-and stereo-selective sites (Experiment 5). **Panel A** shows the level of sensitivity to direction of motion (centrifugal vs. centripetal) across different ROIs. **Panel B** shows the sensitivity of response to the stimulus depth (near vs. far). Motion and stereo-selective sites show stronger sensitivity to change in the stimulus motion direction and depth, respectively. Error bars show one standard error of mean.

By the same token, stereo-selective sites are expected to be more selectively involved in discriminating stimuli based on their stereoscopic depth ^3, 52-54^. To compare the level of depth sensitivity between motion-and stereo-selective sites, a subset of subjects (*n=*11; Table S1) were presented with RDS (with 0° – 0.2° binocular disparity), oscillating either in front (i.e. nearer to the observer) or behind the fixation fronto-parallel plane (i.e. farther) (see Online Methods). Depth sensitivity was defined as the absolute difference between the responses evoked by nearer vs. farther depths. As presented in Figure 5B, we found a significantly stronger depth sensitivity within stereo-compared to the motion-selective sites (*p*<0.01; see also Table S10). Here, the overall level of depth sensitivity remained mostly the same between V2, V3, and V3A (*p*=0.63). These results support the hypothesis that, compared to motion-selective sites, stereo-selective sites contribute more strongly in signaling depth variation.

### Experiment 6. Selective functional connectivity between functionally alike versus unalike sites

Lastly, we tested whether motion-and stereo-selective sites in V2, V3 and V3A, showed stronger functional connections to sites with the same (‘alike’) rather than different (‘unalike’) selectivity (Figure 6A). If confirmed, this would support the hypothesis that motion-and stereo-selective sites comprise two channels within the magnocellular stream to process motion and stereopsis.

**Figure 6.**
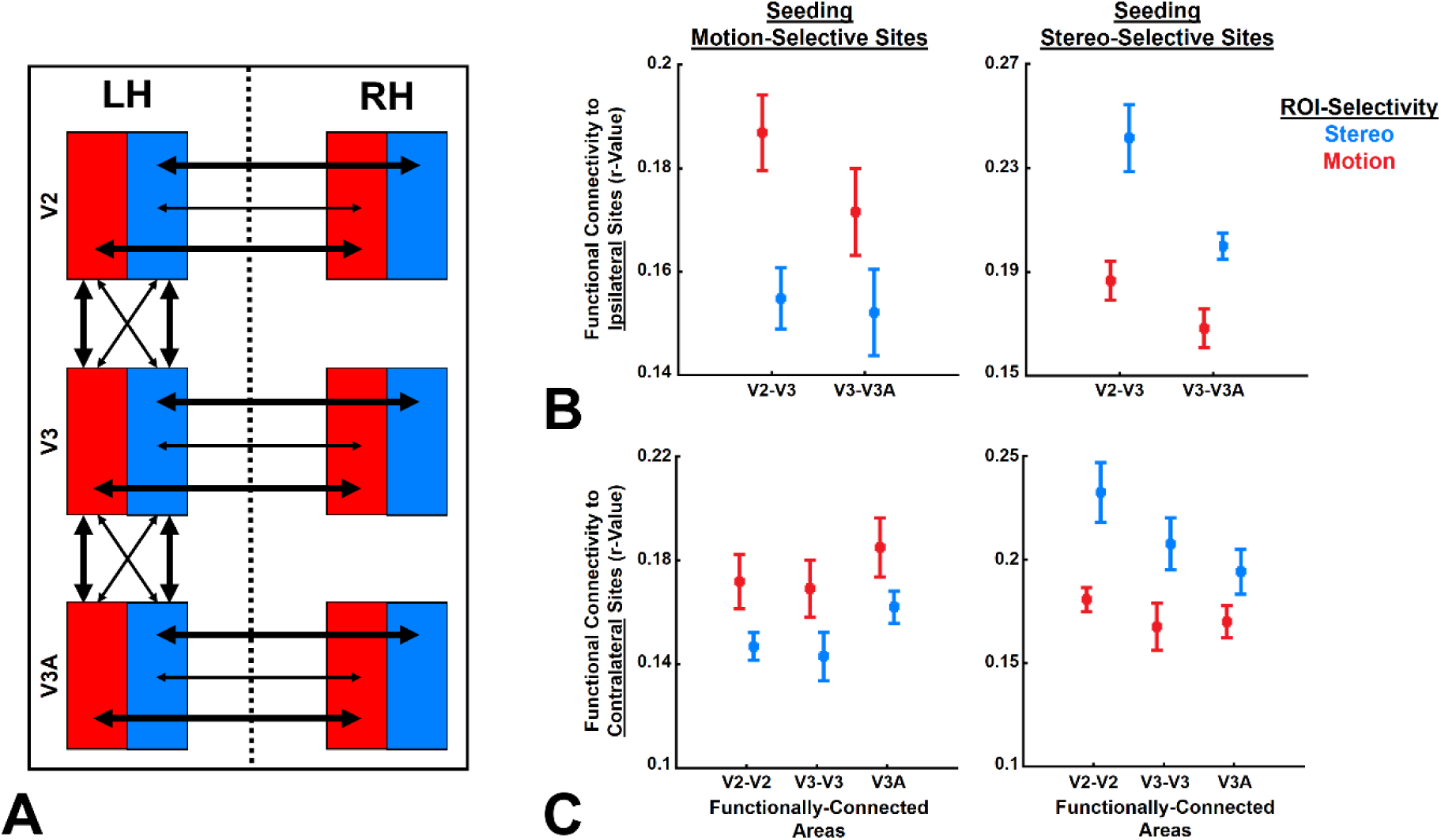
The functional connection between ipsilateral and contralateral ROIs measured during resting state with closed eyes (Experiment 6). **Panels A** shows a schematic representation of the proposed hypothesis about the functional connection between the sites. Motion-and stereo-selective sites are depicted as red and blue boxes. Stronger/weaker functional connections are also depicted with thick/thin arrows. **Panel B** shows the level of functional connectivity between V2-V3 and V3-V3A. Here, the seeded areas and the ROIs were located within the same hemisphere. **Panel C** shows the level of functional connectivity between the two hemispheres (i.e. the seeded area and the ROI were located contralaterally). In **Panels B and C**, the left vs. right column show the results of seeding motion vs. stereo-selective sites. In each hemisphere, motion-and stereo-selective sites show stronger functional connectivity to ipsilateral and contralateral sites with the same rather than different selectivity. Error bars show one standard error of mean.

To test this, we used the method of resting-state functional connectivity (see Online Methods). Based on an independent set of scans, and in the absence of any visual input (eyes closed), we measured the level of co-fluctuations between spontaneous activity evoked within motion-and stereo-selective sites. The results showed that (Figures 6B), in adjacent areas, functional connections were significantly stronger between alike (i.e. motion-to-motion and stereo-to-stereo) rather than unalike sites (i.e. motion-to-stereo) (*p*<0.01; see also Table S11). A similar result was found when we tested the functional connection between contralateral hemispheres (Figure 6C). The latter result ruled out the possibility that the selective functional connectivity between alike sites could be due to the smaller distance between them, compared to unalike sites.

Notably, this result could not be attributed to a common sensory input, because all recordings were conducted during the resting state with eyes closed. Together, these results supported the hypothesis that motion-and stereo-selective sites comprise two interdigitated fine-scale channels within the magnocellular stream.

## Discussion

For decades, the concept of segregated magnocellular vs. parvocellular pathways shaped our understanding of distributed neuronal processing across visual areas ^1, 2, 10, 11^. Here we provide direct evidence for the existence of two mesoscale processing channels within the magnocellular stream that selectively contribute to motion and stereopsis encoding. As we discuss below, the existence of such fine-scale channels in human extrastriate visual cortex was completely unknown to us and could not be predicted based on animal models. This is an important milestone for studying human brain function, especially in those regions that are likely different between humans and NHPs.

### Consistent but beyond animal models for area V2

Evidence for segregated magnocellular vs. parvocellular streams in area V2 of NHPs has been provided in multiple studies. Specifically, motion-^4, 51, 55^ and stereo-selective sites ^3, 6, 7^ that comprise the magnocellular stream were localized mainly within thick stripes (see also ^5, 8-10^. These sites also showed strong orientation selectivity ^42, 45^ and weak-to-no color-selective response ^43, 56^. However, none of these studies tested for the relative organization of motion-vs. stereo-selective sites.

Consistent with these studies in NHPs, Experiment 2 showed that motion-and stereo-selective sites in human V2 showed little overlap with V2 color-selective sites. We also found a stronger orientation-selective response in motion-and stereo-compared to color-selective sites. Moreover, we showed evidence for differential SF preference in motion/stereo-vs. color-selective sites as expected from the influence of magnocellular vs. parvocellular streams ^46, 47^. Thus, our findings regarding the interdigitation of motion/stereo-vs. color-selective sites, plus a stronger influence of the magnocellular stream on the function of motion/stereo-selective sites are consistent with previous reports in NHPs (see above).

However, the evidence provided here for the interdigitated organization of motion-and stereo-selective sites across human visual areas V2, V3 and V3A, could not be predicted from previous studies in NHPs. This is mainly because methodological limitations prohibited localizing motion-and stereo-selective sites within the same animal. In this condition, and regarding the between-subject-variability in the organization of motion-and stereo-selective sites (also see Figures 2 and 3), it is impossible to examine the level of overlap between these sites when they were mapped in different animals. This said, our current findings do not rule out the possibility that a homologous organization may also exist in NHPs. But, in absence of such evidence the relative organization of these sites in NHP remains unclear.

### Limited knowledge of area V3 in NHP

Our knowledge of area V3 in NHPs is relatively limited. Although many studies have reported color-, motion-, stereo-and orientation-selective sites within this area ^12-14, 19, 42, 43^, the relative organization of these sites is still a matter of debate. For instance, a recent study suggested a segregated organization of color-and orientation-selective sites within areas V3 and V4 ^44^. While earlier studies reported a significant overlap between these sites within area V3 ^12, 19^. Considering this, plus the smaller size of area V3 in NHPs compared to humans ^57, 58^, it would have been challenging to predict our current findings from the animal model.

### V3A in humans vs. NHP

Multiple studies have reported stereo-selective activity in area V3A of humans ^13, 28^-^30, 52, 59^ and monkeys ^13, 14, 60^. In both species, color-selective sites are rarely found in this region ^2, 40, 43^. However, V3A contribution to motion encoding may differ between humans and NHPs. Specifically, a group of neuroimaging studies have suggested that motion-selective activity in V3A is limited to humans ^23, 24^. While others, based on using single-cell recording, have shown evidence for motion-selective neurons in this area ^20-22^. Thus, interdigitated organization of motion-and stereo-selective sites in area V3A (Figure 3) may (or may not) be limited to humans.

### Selective functional connectivity between alike sites

Previously we showed evidence for selective functional connectivity between “stereo-vs. color-selective” ^28^ and “motion-vs. color-selective” ^31^ sites. Here, we extend those findings by showing that such a selective functional connection also exists between “motion-vs. stereo-selective” sites (Figure 6). Obviously, in absence of any evidence for interdigitated motion-and stereo-selective sites in NHPs (as mentioned above), the selective functional connection between these sites could not be predicted based on previous studies in NHPs.

This selective functional connectivity between alike sites supports the existence of two fine-scale channels within the magnocellular stream. However, this finding does not suggest that these two channels are segregated from each other. Rather it suggests that activity in one site has a greater influence on activity within sites with the same (alike) rather than different (unalike) selectivity.

### Interdigitated but not necessarily independent

It is widely known that stimulus motion may influence depth processing ^48^ and that motion parallax may be used as an independent cue in depth perception ^61^. These behavioral phenomenon are not in contrast to our findings. To clarify, as mentioned above, our findings did *not* suggest that motion-and stereo-selective channels are segregated (independent) from each other. Rather, they indicated that these two channels consisted of interdigitated cortical sites. The exact mechanisms of interaction between these sites, and also between them and those sites that comprise the parvocellular stream, are beyond the focus of this study and needs to be assessed in the future (also see ^62^).

### Limitations

For majority of our subjects, our scan coverage did not include the anterior portion of area V4. For those in whom we had access to area V4 (Figures 1-3), we noticed the same functional organization (i.e. interdigitated color-, motion-and stereo-selective sites) as we detected in the earlier areas. Also, in those individuals for whom we had access to the posterior tip of medial temporal sulcus, we could localize area MT based on its selectivity for motion and the absence of selectivity for color ^34, 36^. Unfortunately, the small number of these cases kept us from expanding our conclusion to those areas.

In almost all fMRI studies, the expansion of sampling area requires either a decrease in the spatial resolution or a decrease in signal-to-noise ratio (e.g. when the method of simultaneous multislice imaging (SMS) ^63^ is used). To achieve this goal without sacrificing the spatial resolution, a combination of stronger MR scanners ^64^, more advanced data acquisition sequences (e.g. ^65^) and more efficient data processing methods (e.g. ^66, 67^) will be required in the future.

Although the current study focused on extrastriate cortex, similar questions could also be explored in the striate visual cortex (V1) and the lateral geniculate nucleus (LGN) of the thalamus. In both regions, magnocellular and parvocellular streams are expected to influence separate sites ^46, 47, 68-71^. However, the spatial resolution used here (1 mm isotropic) was inadequate to differentiate these sites and to examine their fine-scale functional organization. Moreover, LGN was located outside of the sampled region in all subjects. Here again, more technological advances are required to grant us access to those regions with adequate spatial resolution (also see ^72, 73^).

## Conclusion

Until recently, applications of fMRI have been mostly limited to localization of large-scale brain areas and assessing their function across various experimental conditions. This has changed with introduction of high-resolution fMRI using ultra-high field scanners (e.g. 7 or 9.4 Tesla). Using advanced technologies and the state-of-the-art scanners, many groups have revealed mesoscale structures within the spatial scale of cortical columns in human striate ^28, 74, 75^ and extrastriate cortex ^28, 30-32^. For understandable reasons, these studies typically limited themselves to replicating what was already known to us based on animal models. However, with improvement in the reliability of fMRI, the present study demonstrates how human fMRI studies can be used to test new hypotheses beyond what was already known to us based on animal models.

## Acknowledgment

This work was supported by NIH NEI (grants R01EY026881 and R01 EY030434), and by the MGH/HST Athinoula A. Martinos Center for Biomedical Imaging. Crucial resources were made available by a NIH Shared Instrumentation Grant S10-RR019371. We thank Ms. Azma Mareyam for help with hardware maintenance during this study and Dr. Anna Blazejewska for her helps with implementing the method of radial smoothing. We also thank Drs. Roger Tootell, Bruce Rosen, Anna Roe, Haidong Lu, Ichiro Fujita, Gregory DeAngelis and Akiyuki Anzai for their helpful comments.

