## Supplementary material for "Two interdigitated fine-scale channels for encoding motion and stereopsis within the human magnocellular stream": Online Methods

**Participants**

Fifteen human subjects (6 females), aged 23–44 years old, participated in this study (see Table S1). All subjects had normal or corrected-to-normal [visual acuity](https://www.sciencedirect.com/topics/medicine-and-dentistry/visual-acuity) (based on the Snellen test), normal color vision (Ishihara and Farnsworth D15 tests), normal stereovision (Randot test), and radiologically normal brains without history of neuropsychological disorder. All experimental procedures conformed to NIH guidelines and were approved by Massachusetts General Hospital protocols. Written informed consent was obtained from all subjects prior to all experiments.

**General Procedures**

Subjects were scanned multiple times (Table S1) in an ultra-high field scanner (Siemens 7T whole-body system, Siemens Healthcare, Erlangen, Germany) for functional experiments. Details of these scans are listed below. All subjects were also scanned in a 3T scanner (Tim Trio, Siemens Healthcare) for structural imaging.

In all functional experiments (except for the functional connectivity test), stimuli were presented via an LCD projector (1024 × 768 pixel resolution, 60 Hz refresh rate) onto a rear-projection screen, viewed through a mirror mounted on the receive coil array. During these experiments, subjects were instructed to look at a centrally presented object (radius = 0.15°) and to do an orthogonal dummy task. The task was irrelevant to the stimuli presented in the background (e.g. shape change detection (square to circle and vice versa) for the fixation object) and remained the same across the whole run, including the blank presentation interval. Matlab 2020a (MathWorks, Natick, MA, USA) and the Psychophysics Toolbox ^1, 2^ were used to control stimulus presentation.

**Experiment 1a – Localizing motion-selective sites**

Motion-selective sites were localized based on their stronger response to moving compared to stationary stimuli. Stimuli consisted of concentric rings, extending 20° × 26° (height × width) in the visual field, presented against a light gray background (Figure 1a). The experiment was block-designed (block duration = 24 s). In half of the blocks, rings moved radially (centrifugally vs. centripetally; 4°/s) and the direction of motion changed every 3 s to reduce the impact of motion after-effects. In the other half of blocks, rings remained stationary during the whole block. Each run started and finished with a 12 s of uniform gray presentation. The sequence of moving and stationary blocks was pseudo-randomized across runs (10 blocks per run) and each subject participate in 6 runs.

**Experiment 1b – Localizing stereo-selective sites**

Stereo-selective sites were localized based on their stronger response to 3D compared to 2D stimuli (block-designed). Stimuli were sparse (5%) random dot stereograms (RDS), based on red or green dots (0.09° × 0.09°), presented against a black background (Figure 1b). Stimuli extended 20° × 26° in the visual field. Subjects viewed the stimulus through custom made anaglyph spectacles mounted to the head coil. In the 3D blocks, the RDS stimuli formed a stereoscopic percept of an array of cuboids that oscillated in depth (-0.22° to 0.22°; 0.3 Hz), with independent phase. In the 2D blocks, the fused percept formed a fronto-parallel plane intersecting the fixation target (i.e. zero depth at that point) and oscillated translationally (left to right and vice versa; 0.3 Hz). Each experimental run began and ended with 12 s of uniform gray (‘blank’) and included 8 stimulus blocks (24 s per block). Each subject participated in 12 runs and the sequence of blocks was pseudo-randomized across runs.

**Experiments 2a and 2b – Localizing color-selective sites and measuring orientation sensitivity**

To localize color-selective sites, subjects were presented in separate blocks (24 s per block) with sinusoidal gratings (0.2 cycle/degree; 20° × 26°) which varied in either color (between red and blue) or achromatic luminance (Figure 1C). For each subject, colors were adjusted to be equal in luminance across all eccentricities stimulated, using the method of flicker-photometry ^3, 4^ and according to each subject’s color perception (for more details see ^5^).

In different blocks, orientation of colorful and achromatic gratings were either 0°, 45°, 90° or 135°, drifting in orthogonal directions (reversed every 6 s) at 4°/s. This design enabled us to measure the orientation sensitivity of the evoked response (Experiment 2b). However, only the response evoked by achromatic stimuli was used to assess the orientation sensitivity. Each run started and finished with a short block (18 s) of uniform gray of equal mean luminance. The sequence of blocks within each run (9 blocks per run) was pseudo-randomized and each subject participated in 12 runs.

**Experiment 2c – Measuring spatial frequency preference**

To measure the spatial frequency preference in different sites, subjects were presented with gratings (20° × 26°) of differing spatial frequency (0.1, 0.27, 0.73, 2.08 and 5.79 cycle/degree) across different blocks (i.e. block-design). In each block, the spatial frequency of the stimuli remained the same. But the orientation of gratings changed pseudo-randomly every 4 seconds. In addition, grating phase reversed every 1 s. Each run included 15 blocks of 16 seconds each, beginning and ending with an additional block (12 s) of uniform gray of equivalent mean luminance. The sequence of blocks was pseudo-randomized across runs and each subject participate in 12 runs.

**Experiment 3a – Measuring motion-selective activity induced by radially-moving random dots**

In this experiment, rather than using concentric rings (used in Experiment 1a), subjects were presented random dots (Figure 4A). In different blocks, dots moved (4°/s) either centrifugally (33% of blocks), centripetally (33% of blocks) or remained stationary (the rest of blocks). This design enabled us to assess sensitivity of the evoked response to motion direction (centrifugal vs. centripetal) (see Results – Experiment 5). Stimuli were presented against a black background, extending 20° × 26° in the visual field. The size of dots increased with eccentricity according to cortical magnification factor ^6^. Each run included 13 blocks of 24 seconds each, beginning and ending with an additional block (12 s) of uniform black. The sequence of blocks was pseudo-randomized across runs and each subject participate in at least 12 runs.

**Experiment 3b – Measuring motion-selective activity induced by translationally- moving random dots**

The stimuli were similar to those used in Experiment 3a with two exceptions. First, rather than moving radially, stimuli moved either left-ward (20% of blocks), right-ward (20% of block), up-ward (20% of block), and down-ward (20% of block) (Figure S3). This design enabled us to measure the sensitivity of evoked response to motion direction (see above and Results – Experiment 5). In the rest of blocks, stimuli remained stationary. Second, the size of dots remained the same (radius = 0.02°) and did not change with their eccentricity. The other aspects of the procedure were similar to those in Experiment 3a.

**Experiment 3c – Measuring stereo-selective activity induced by depth-varying gratings**

In this experiment, rather than using RDS to measure stereo-selective activation, (used in Experiment 1b), subjects were presented with low SF (0.25 cycle/degree) red and green gratings (Figure 4C). Subjects viewed the stimuli through custom anaglyph spectacles as in Experiment 1b. The experiment was block-designed and, stimuli overlaid and fused within all experiment blocks. In 50% of blocks, the level of binocular disparity oscillated between -0.22° to 0.22° (0.3 Hz), forming a stereoscopic percept of oscillation in depth. In the rest of blocks, the level of binocular disparity remained equal to zero and stimuli formed a perception of oscillation in the fronto-parallel plane. Stimuli were presented against a black background, extending 20° x 26° in the visual field. The orientation of gratings varied randomly between blocks. The other aspects of the procedure was similar to Experiment 1b.

**Experiment 4 – Measuring stereo-selective activity induced by 3D natural scenes**

Subjects were presented with 3D vs. 2D scenes (Figure S6) across different blocks. Stimuli included pictures of indoor and outdoor scenes, selected from Southampton-York Natural Scenes (SYNS) dataset ^7^. Each stimulus extending 20° × 26° of visual field. Subjects viewed the stimuli through custom anaglyph spectacles as in Experiments 1b and 3c. For 3D stimuli, the level of binocular disparity varied between -0.15° to 0.15° in different parts of each image (Figure S6). Each block contained 24 stimuli (1 s per stimuli) with no blank presentation between the stimuli. The sequence of stimuli within the blocks was randomized. Each subject participated in 12 runs (11 blocks per run), beginning and ending with an additional block (12 s) of uniform black presentation.

**Experiment 5 – Measuring depth sensitivity**

To measure the depth sensitivity of the response evoked within stereo- and motion-selective sites, subjects were presented with RDS stimuli (20° × 26°) similar to those used in Experiment 1b. But here, in different blocks, stimuli were oscillating (0-0.22°) either ‘in front’ or ‘behind’ a fronto-parallel plane that intersected the fixation target. Each experimental run included 9 stimulus blocks (24 s per block) and each run began and ended with control conditions of 12 s of uniform gray (‘blank’). Other details of the stimuli are similar to those in Experiment 1b. Also notably, direction-sensitivity was measured based on the data collected during Experiments 3a and 3b.

**Experiment 6 – Resting-state functional connectivity**

In these scans, subjects were instructed to keep their eyes closed during the whole scan, but not to sleep. Each scan session consisted of 6 runs, and each run took 256 s. Experimenters talked to the subject between each run to ensure wakefulness.

**Retinotopic Mapping**

For all subjects the border of retinotopic areas and the presentation of central (radius = 0-3°) and peripheral (radius = 3-10°) visual fields were defined retinotopically ^6^. Stimuli were based on a flashing radial checkerboard, presented within retinotopically limited apertures, against a gray background. These retinotopic apertures included wedge-shaped apertures radially centered along the horizontal and vertical meridians (polar angle = 30°), plus a central disk (radius = 0-3°) and a peripheral ring (radius = 3-10°). These stimuli were presented to subjects in different blocks (24 s). The sequence of blocks was pseudo-randomized across runs (8 blocks per run) and each subject participate in at least 4 runs.

**Imaging**

Functional experiments (see above) were conducted in a 7T Siemens whole-body scanner equipped with SC72 body gradients (70 mT/m maximum gradient strength and 200 T/m/s maximum slew rate) using a custom-built 32-channel helmet receive coil array and a birdcage volume transmit coil. Voxel dimensions were nominally 1.0 mm. We used single-shot gradient-echo EPI to acquire functional images with the following protocol parameter values: TR=3000 ms, TE=28 ms, flip angle=78°, matrix=192×192, BW=1184 Hz/pix, echo-spacing=1 ms, 7/8 phase partial Fourier, FOV=192×192 mm, 44 oblique-coronal slices, acceleration factor *R*=4 with GRAPPA reconstruction and FLEET-ACS data ^8^ with 10° flip angle. The field of view included occipital cortical areas V1, V2, V3 and the posterior parts of V4v and V4d.

Structural (anatomical) data were acquired using a 3D T1-weighted MPRAGE sequence with protocol parameter values: TR=2530 ms, TE=3.39 ms, TI=1100 ms, flip angle=7°, BW=200 Hz/pix, echo spacing=8.2 ms, voxel size=1.0 × 1.0 × 1.33 mm^3^, FOV=256 × 256 × 170 mm^3^.

**General data analysis**

Functional and anatomical MRI data were pre-processed and analyzed using FreeSurfer and FS-FAST (version 6.0; <http://surfer.nmr.mgh.harvard.edu/>) ^9^. For each subject, inflated and flattened cortical surfaces were reconstructed based on the high-resolution anatomical data ^10-12^. To enable intra-cortical smoothing (see below), we also automatically generated, in addition to the standard pial surface (i.e., the gray matter border with the surrounding cerebrospinal fluid or CSF) and the white matter (WM) surface reconstructions, a family of 11 intermediated equidistant surfaces spaced at intervals of 10% of the cortical thickness, which consisted the WM-GM (surface 0) interface surface, the GM-CSF (surface 10) interface surface, and 9 intermediate surfaces within gray matter.

All functional images were corrected for motion artifacts. For each subject, functional data from each run were rigidly aligned (6 DOF) relative to their own structural scan using rigid Boundary-Based Registration ^13^. This procedure enabled us to average data collected for each subject across multiple scan sessions.

No *tangential* spatial smoothing was applied to the main imaging data acquired at 7T (i.e. 0 mm FWHM). Rather we used the more advanced method of radial (intracortical) smoothing (Blazejewska et al., 2019) – i.e. perpendicular to the cortex and within the cortical columns. The extent of this radial smoothing was limited to the bottom 30% of the gray-matter thickness starting from the gray-white matter interface. This approach allowed us to retain spatial resolution, while increasing SNR through exploiting the prior knowledge about the columnar organization within the regions of interest ^5, 14, 15^. Moreover, by not sampling from more superficial layers, we avoid spatial blurring caused by large veins at the pial surface ^5, 16-18^.

A standard hemodynamic model based on a gamma function was fit to the fMRI signal to estimate the amplitude of the BOLD response. For each individual subject, the average BOLD response maps were calculated for each condition ^19^. Finally, voxel-wise statistical tests were conducted by computing contrasts based on a univariate general linear model, and the resultant significance maps were projected onto the subject’s anatomical volumes and reconstructed cortical surfaces.

**Analysis of overlap**

To measure the level of overlap between sites, the selectivity maps (e.g. Figure 1A-C) were first thresholded at *p*<0.05. Then, all thresholded values in areas V2, V3 and V3A were normalized (min-to-max were converted linearly to 0-to-1). A site was called selective if the resultant normalized value for one feature (i.e. either motion, stereopsis or color) was $\sqrt{2}$ time larger than that the value measured for the others.

To test whether the selectivity maps were overlapping or not, we also measured the extent of overlap when the organization of motion-, stereo- and color-selective maps vertices was spatially ‘shuffled’. For each subject, this shuffling process was repeated 10,000 times and the averaged level of overlap was used as the chance level (for that individual). If two selective sites were overlapping, we expected their level overlap to be significantly above the chance level.

**Functional connectivity analysis**

Details of the functional connectivity analysis are similar to those reported previously ^5, 14, 20^. Briefly, after preprocessing (see above), for each subject we removed sources of variance of non-interest including: all motion parameters measured during the motion correction procedure, the global signal, the mean signal from the portion of ventricles that were included in the acquired EPI slices, and the mean signal from a region within the deep cerebral white matter. Then, we extracted the mean BOLD signal time course for V2/V3/V3A motion- and stereo-selective sites to use as seeds in a seed-based connectivity analysis. The correlation coefficient was computed for each of these time course seeds against the preprocessed resting-state time course data, from every voxel from the ipsilateral and contralateral hemisphere.

**Region of interest (ROI) analysis**

ROIs including motion-, stereo- and color-selective sites across areas V2, V3 and V3A, defined for each subject based on their own data (Experiments 1 and 2) and retinotopy mapping (see above). To remove the impact of the fixation object presentation and the corresponding task as well as adjust the eccentricity of ROIs, only sites across 3–10° eccentricities were used in ROI analysis. Sites that showed overlapping selectivity were excluded from the ROIs. To improve sensitivity in all analyses, data from the left and right hemispheres were averaged.

**Statistical data analysis**

To examine the significance of independent parameters in each experiment, we used either paired t-test or repeated-measures ANOVA. Repeated-measures ANOVA is particularly susceptible to the violation of sphericity assumption, caused by the correlation between measured values and unequal variance of differences between experimental conditions. To address this problem, when necessary (determined using a Mauchly test), results were corrected for violation of the sphericity assumption, using the Greenhouse-Geisser method.

**Data availability statement**

Data will be shared upon request.

**Studies Cited in the Online Methods**
