## Supplementary material for "Two interdigitated fine-scale channels for encoding motion and stereopsis within the human magnocellular stream": Supp. Figures

**Supplementary Figures**

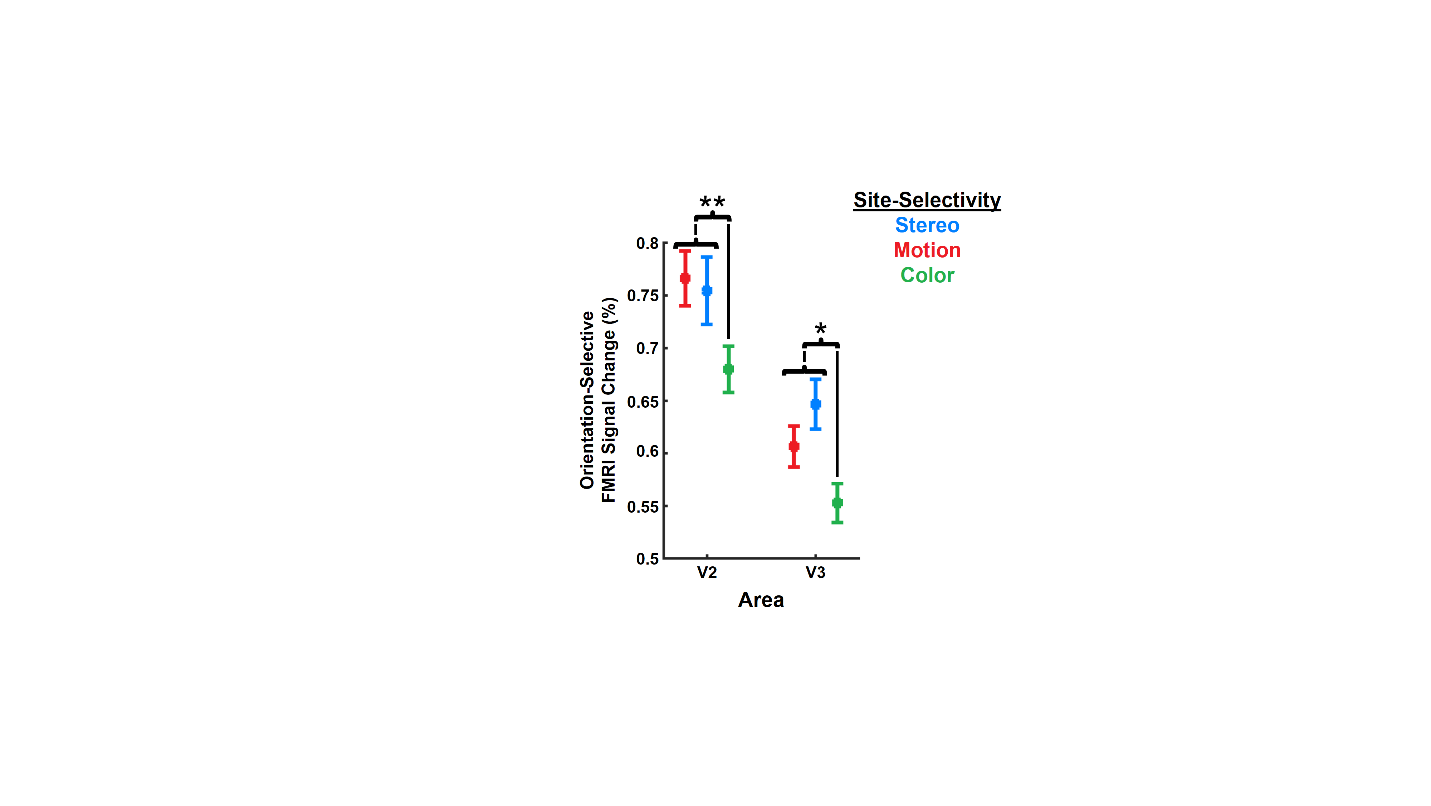

**Figure S1)** The level of orientation selectivity measured across stereo-, motion- and color-selective sites within areas V2 and V3 (Experiment 2b). Consistent with previous findings in NHPs, the level of orientation-sensitivity is higher stereo- and motion-selective sites compared to color-selective sites. The difference between motion- vs. stereo-selective sites was comparably small. Area V3A is excluded from this analysis because color-selective sites are rarely detected within this area. Error bars show one standard error of mean. (**: *p*<0.01 and *: *p*<0.05)

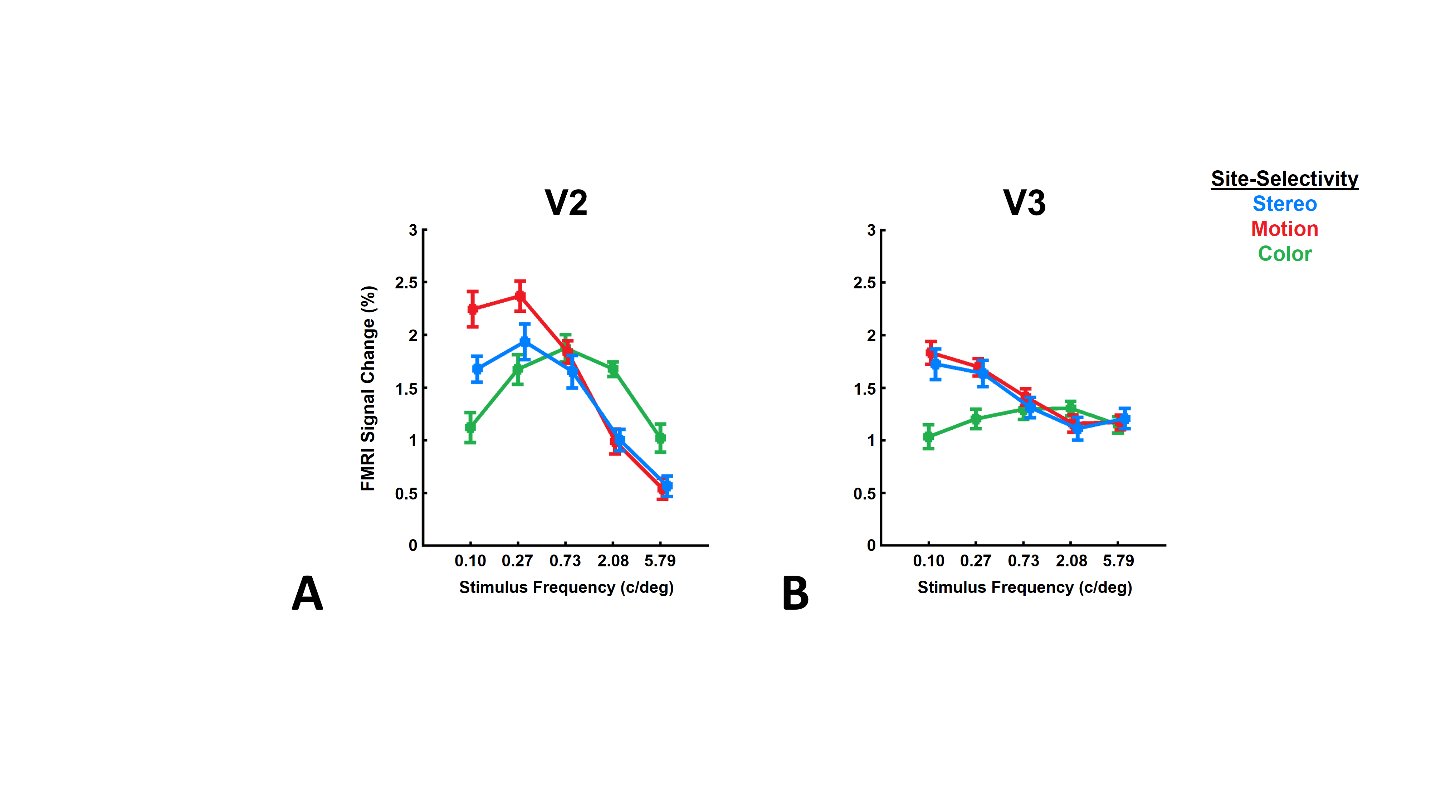

**Figure S2)** SF preference measured across stereo-, motion- and color-selective sites within areas V2 and V3 (Experiment 2c). **Panels A and B** show the activity measured in areas V2 and V3, respectively. In both areas, stereo- and motion-selective areas showed a preference to lower SFs. Comparably, color-selective sites showed preference for higher-SFs. These findings are consistent with the hypothesis that the magnocellular stream has a stronger influence on the activity evoked within motion- and stereo- compared to color-selective sites. Error bars show one standard error of mean.

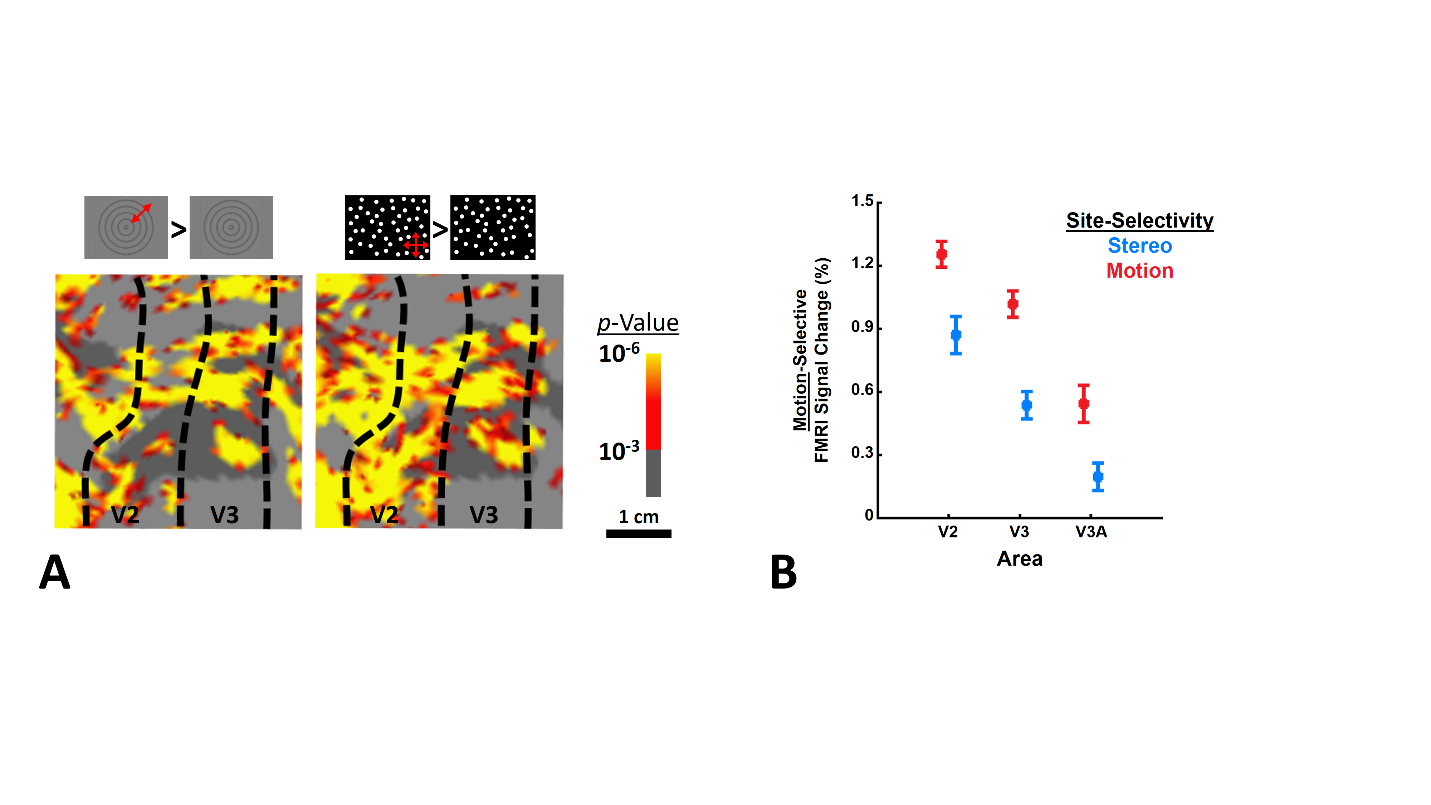

**Figure S3)** The organization of motion-selective sites evoked by translationally moving vs. stationary random dots (Experiment 3b). **Panel A** shows the location of motion-selective sites in one individual (other the one used in Figures 1-3) based on moving vs. stationary concentric rings (left) and translationally moving vs. stationary random-dots (right). **Panel B** shows the level of motion-selective activity across areas V2, V3 and V3A, evoked by translationally moving (vs. stationary) stimuli. The regions of interest in each area were localized independently based on concentric rings (Experiment 1b). Consistent with activity maps and **Figure 4B**, the level of selective activity evoked by moving random-dots was higher in motion- compared to stereo-selective sites. Error bars show one standard error of mean.

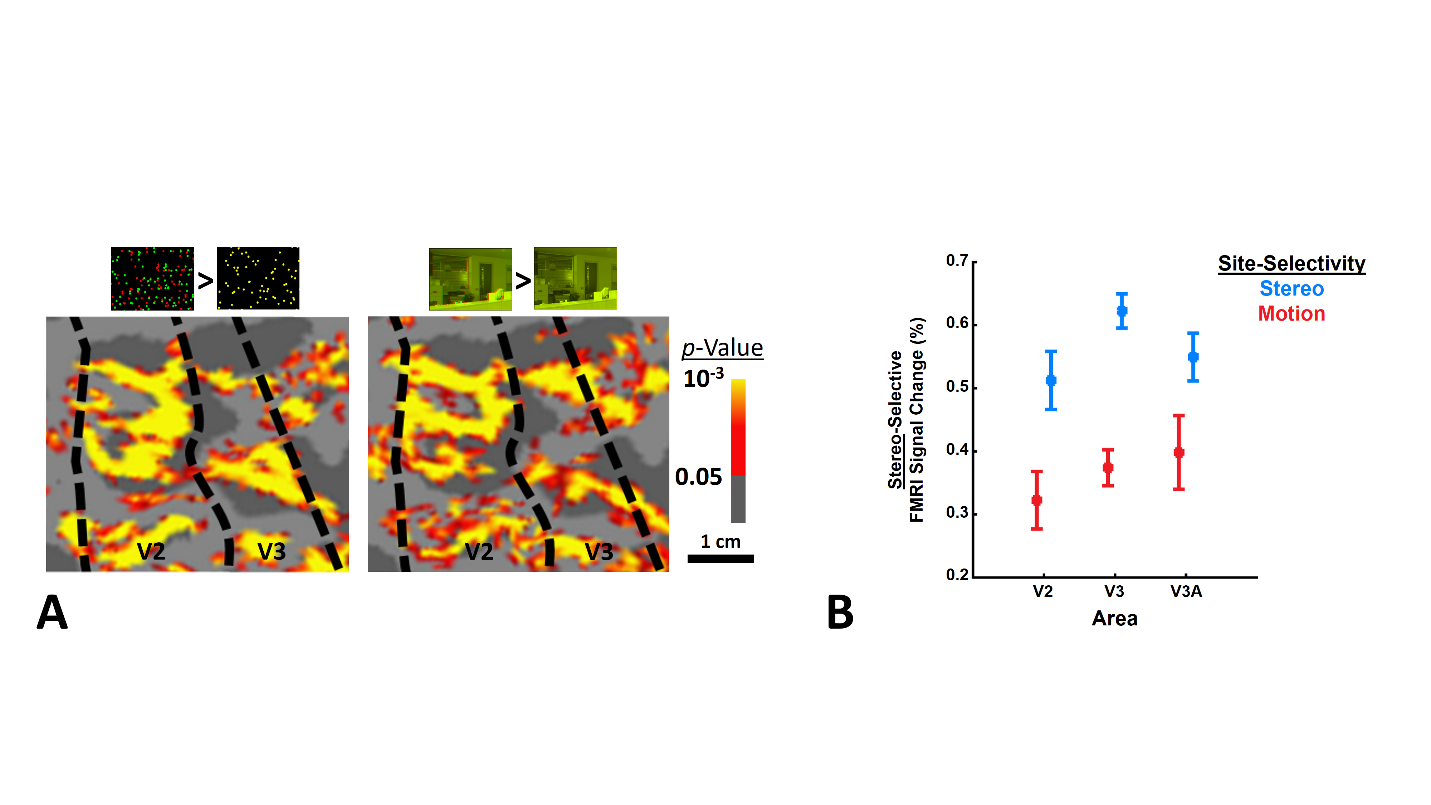

**Figure S4)** Localization of stereo-selective sites in absence of binocular disparity oscillation (Experiment 4). **Panel A** shows the location of stereo-selective sites in one individual (other the one used in Figures 1-3) based on oscillating RDS (left) and non-oscillating scenes (right; see also Figure S6). The overall organization of stereo-selective sites remained mostly unchanged in absence of disparity oscillation. **Panel B** shows the level of stereo-selective activity across areas V2, V3 and V3A, evoked by 3D vs. 2D scenes. The regions of interest in each area were localized independently based on RDS. Consistent with the activity maps, the level of evoked activity by 3D vs. 2D scenes was higher in stereo- compared to motion-selective sites. Error bars show one standard error of mean.

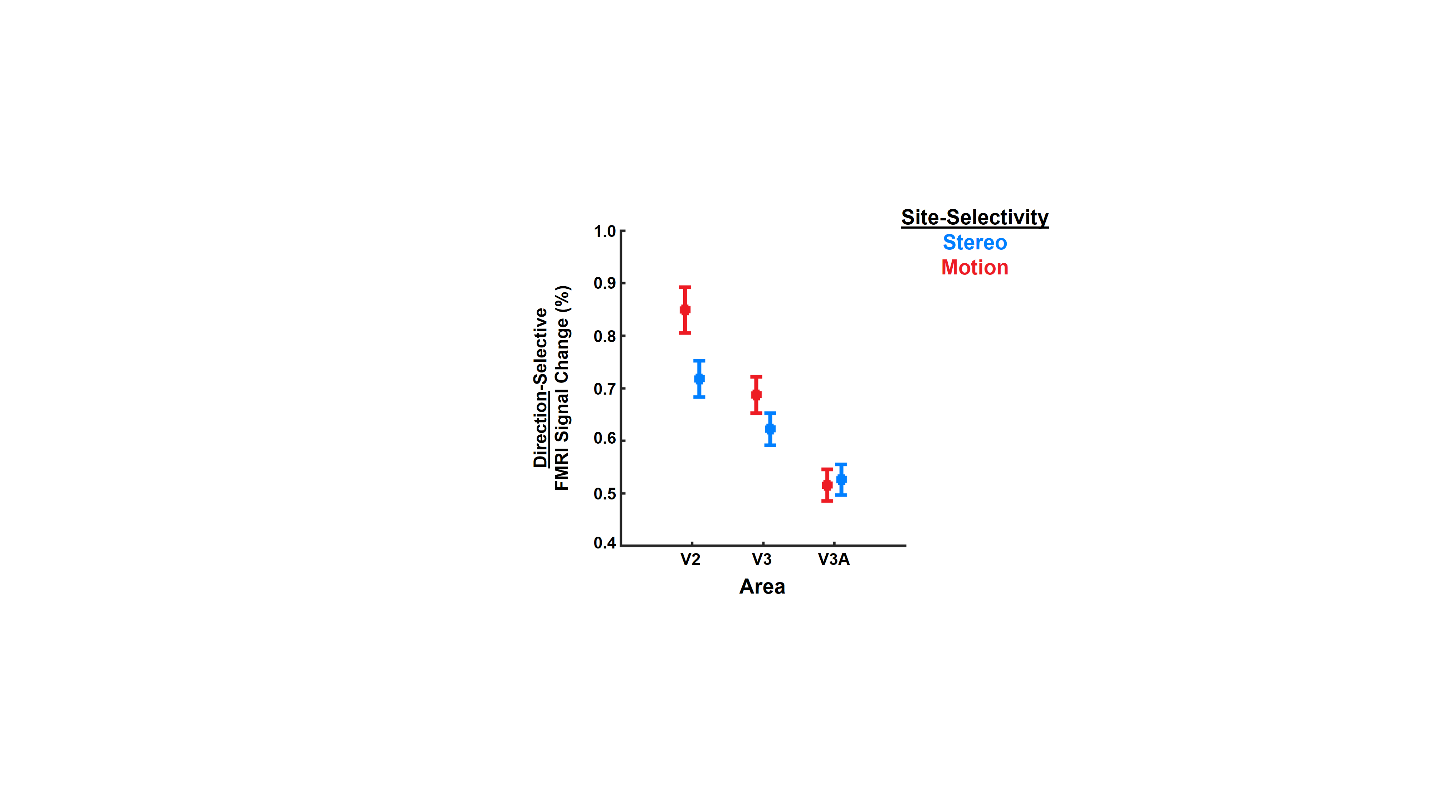

**Figure S5)** Sensitivity to translational motion direction measured in motion- and stereo-selective sites (Experiment 5). Motion-selective (compared to stereo-selective) sites show stronger sensitivity to motion direction. The overall level of direction-sensitivity becomes weaker in V3A compared to V2, most likely due to the simple form of stimuli. Error bars show one standard error of mean.

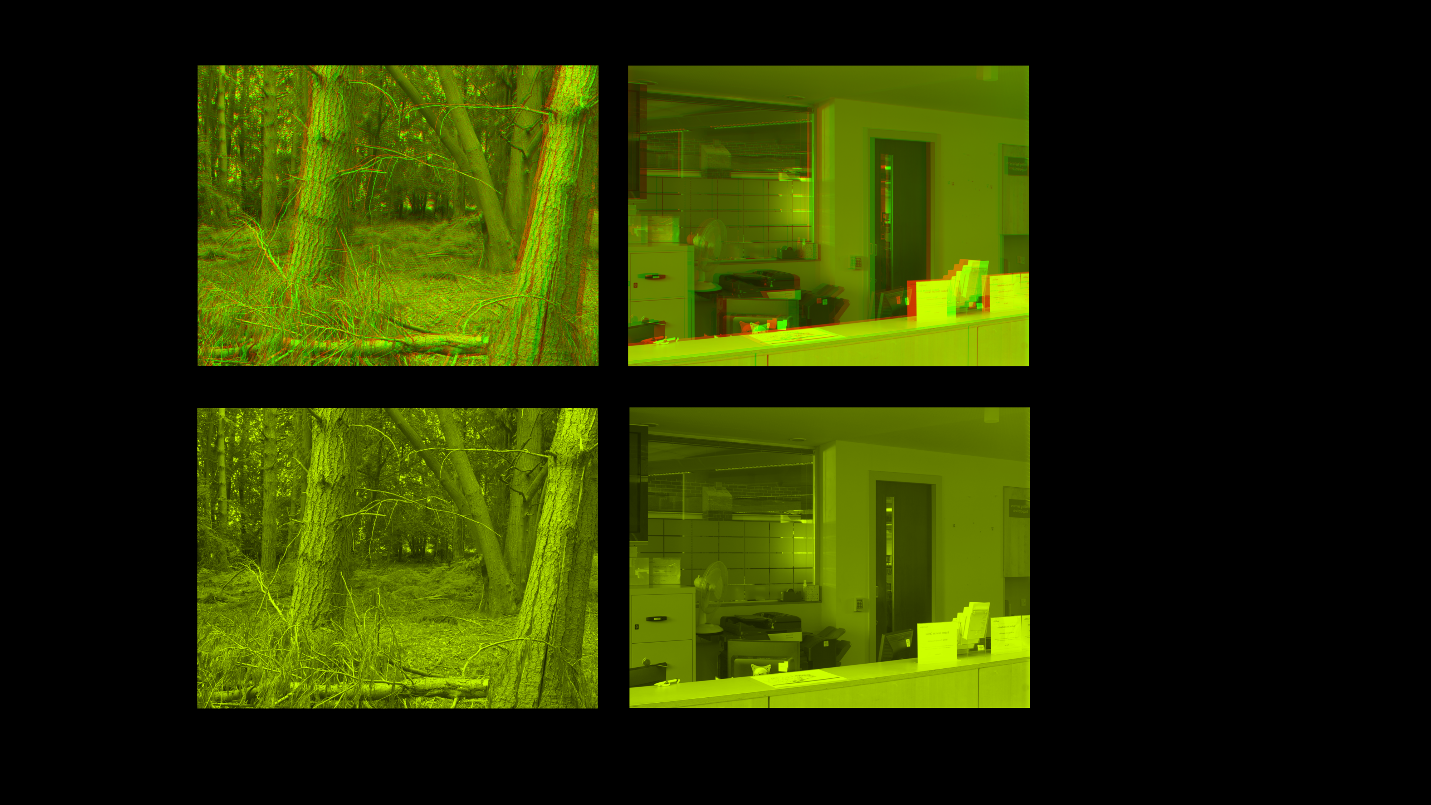

**Figure S6)** Example of stimuli used in Experiment 4. Stimuli included images of outdoor and indoor scenes in 3D (top) or 2D (bottom) formats. Subjects viewed stimuli through anaglyphic spectacles.
