## Supplementary material for "Two interdigitated fine-scale channels for encoding motion and stereopsis within the human magnocellular stream": Supp Tables

**Supplementary Tables**

**Table S1** – Participant demographics

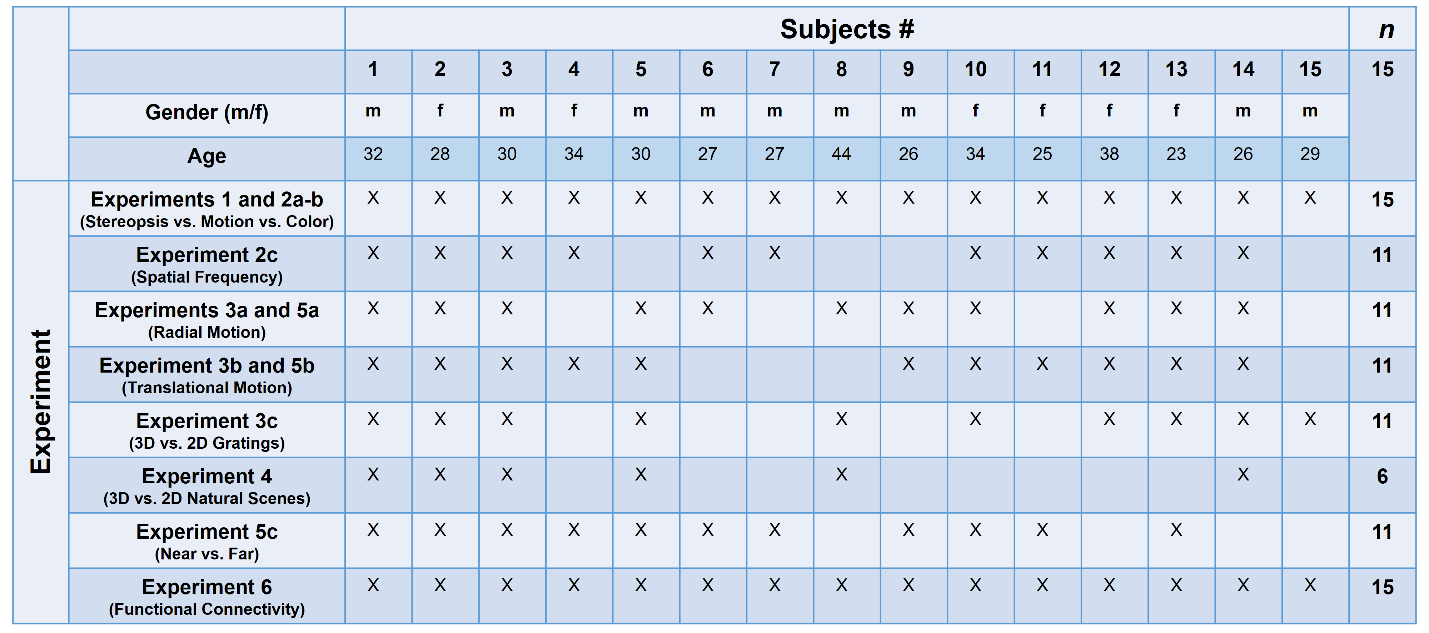

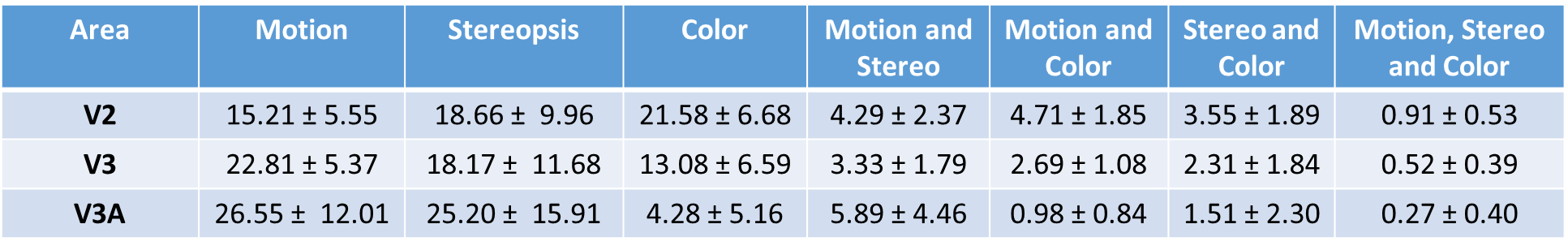

- All sizes are normalized relative to the size of visual area.

**Table S2** – The size of motion-, stereo- and color-selective sites and their level of overlap

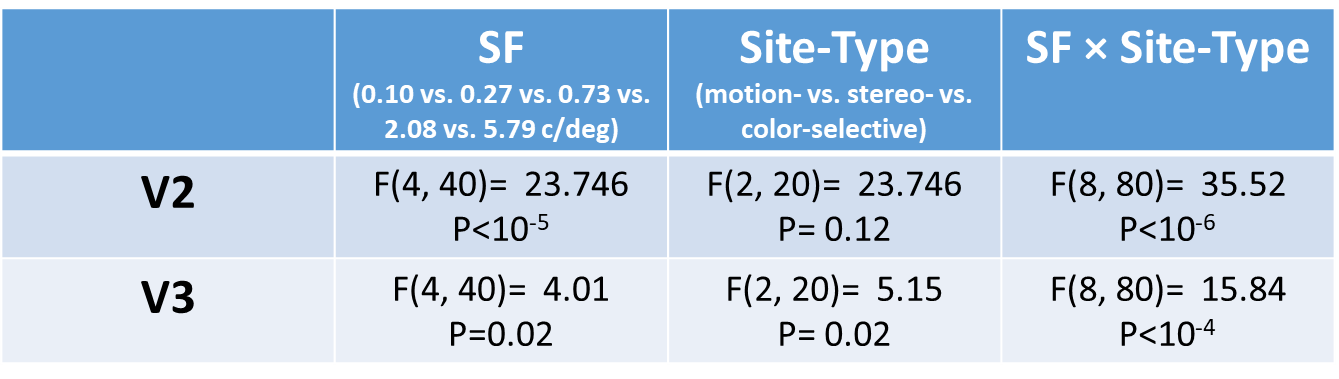

- Results yielded from two separate applications of two-way repeated measures ANOVA to the activity evoked within areas V2 and V3.
- Similar results were found when we applied a single application of three-way ANOVA after including area (V2 vs. V3) as an independent factor.

**Table S3** – In areas V2 and V3, SF preference differs between sites that comprised magnocellular (i.e. motion- and stereo-selective sites) vs. parvocellular (i.e. color-selective sites) streams.

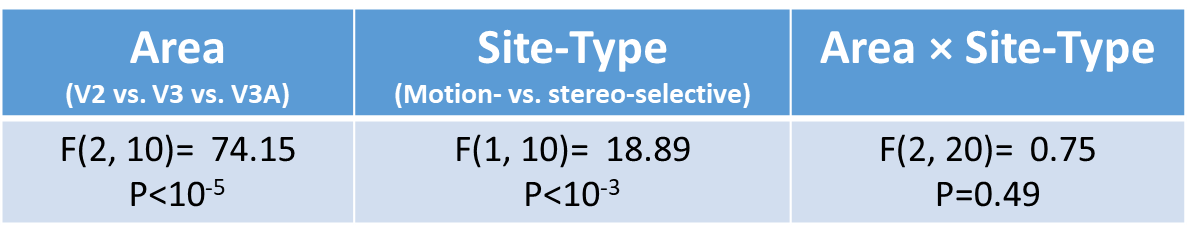

- Results yielded from an application of two-way repeated measures ANOVA to the activity evoked by contrasting the response to moving – stationary stimuli.

**Table S4** – In areas V2, V3 and V3A, radially moving random dots evoked a larger selective response in motion- vs. stereo-selective sites.

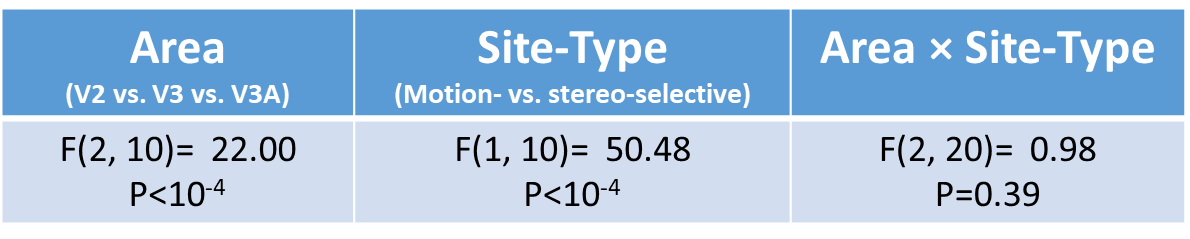

- Results yielded from an application of two-way repeated measures ANOVA to the activity evoked by contrasting the response to moving – stationary stimuli.

**Table S5** – In areas V2, V3 and V3A, translationally moving random dots evoked a larger selective response in motion- vs. stereo-selective sites.

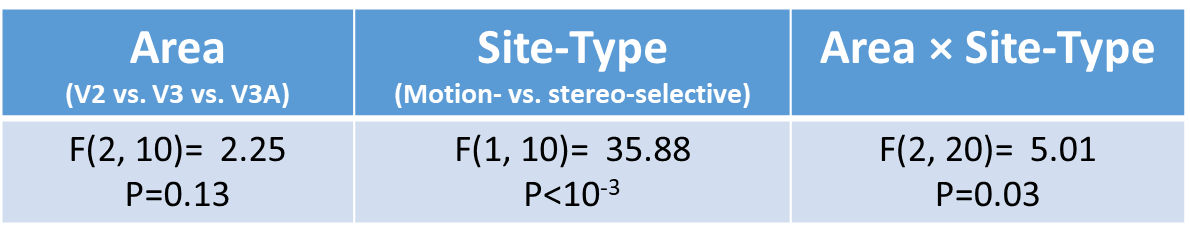

- Results yielded from an application of two-way repeated measures ANOVA to the activity evoked by contrasting the response to stimuli with varying depth (3D) – non-varying (2D).

**Table S6** – In areas V2, V3 and V3A, depth-varying gratings evoked a larger selective response in stereo- vs. motion-selective sites.

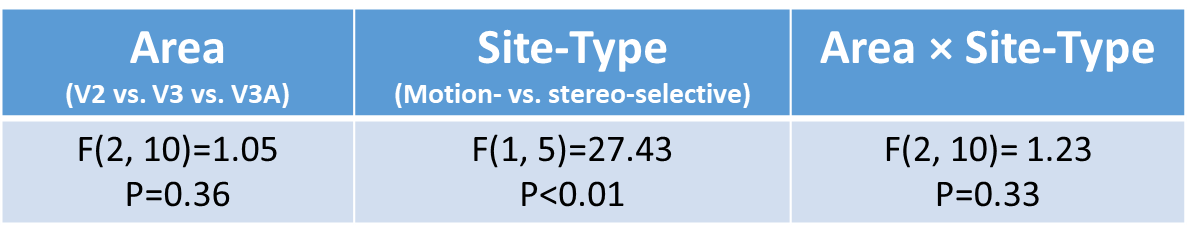

- Results yielded from an application of two-way repeated measures ANOVA to the activity evoked by contrasting the response to 3D – 2D stimuli.

**Table S7** – In areas V2, V3 and V3A, natural scenes with binocular disparity evoked a larger selective response in stereo- vs. motion-selective sites.

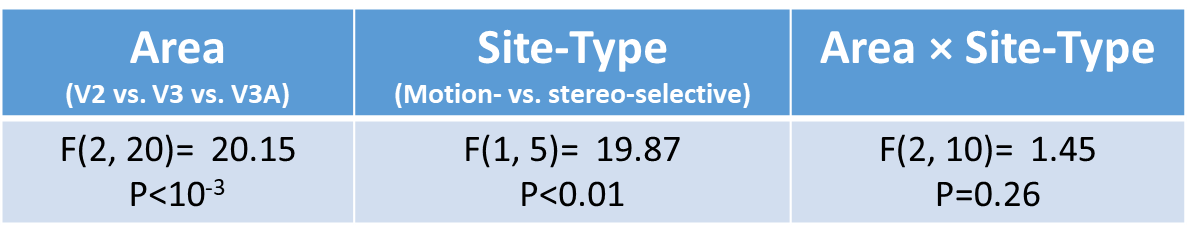

- Results yielded from an application of two-way repeated measures ANOVA to the absolute activity evoked by contrasting the response to centrifugal vs. centripetal motions.

**Table S8** – Radial direction sensitivity is generally stronger in motion- compared to stereo-selective sites.

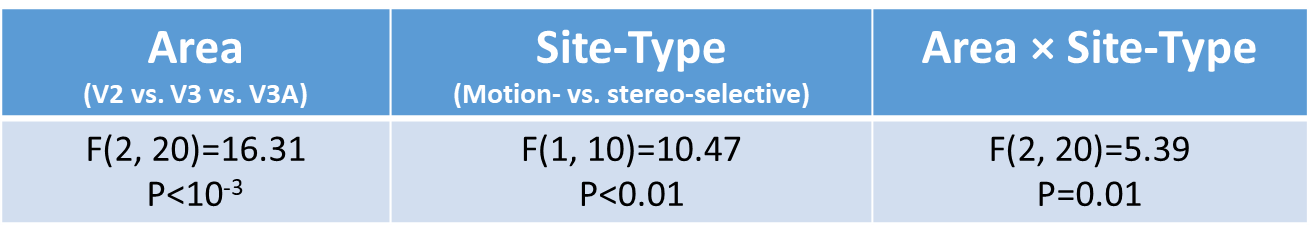

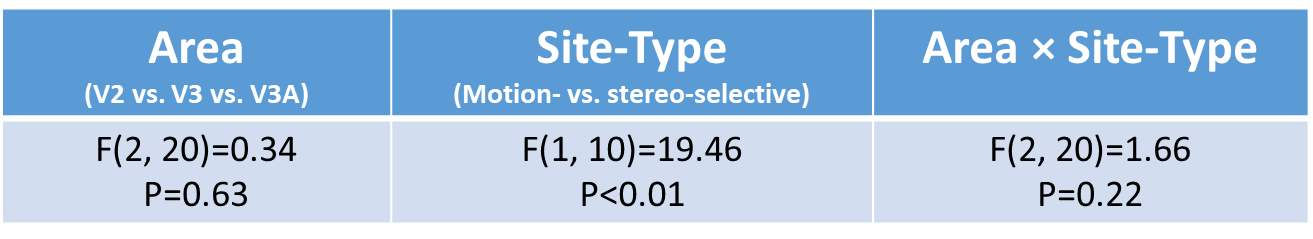

- Results yielded from an application of two-way repeated measures ANOVA to the absolute activity evoked by contrasting the response to different translational motions (upward vs. downward vs. leftward vs. rightward).

**Table S9** – Translational direction sensitivity is generally stronger in motion- compared to stereo-selective sites.

- Results yielded from an application of two-way repeated measures ANOVA to the absolute activity evoked by contrasting the response to RDS stimuli that appeared in front (i.e. nearer) vs. behind (i.e. farther) the fixation object.

**Table S10** – Depth sensitivity is generally stronger in stereo- compared to motion-selective sites.

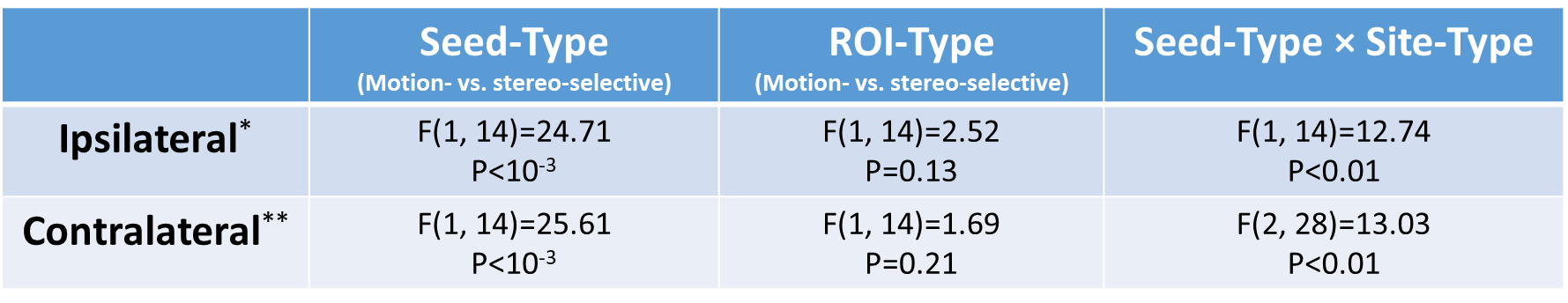

* Results yielded from an applications of three-way repeated measures ANOVA (Area (V2-V3 vs. V3-V3A), seed-type (motion- vs. stereo-selective), and ROI-type (motion- vs. stereo-selective)) to the measured functional connectivity between motion- and stereo-selective sites. Seed sites were located within the ipsilateral hemispheres compared to the ROIs.

** Results yielded from an applications of three-way repeated measures ANOVA (Area (V2 vs. V3 vs. V3A), seed-type (motion- vs. stereo-selective), and ROI-type (motion- vs. stereo-selective)) to the measured functional connectivity between motion- and stereo-selective sites. Seeded sites were located within the contralateral hemispheres compared to the ROIs.

**Table S11** – Functional connectivity is generally stronger between alike compared to unalike sites.
